# Spatial niche separation of an invasive and a native mesopredator

**DOI:** 10.1101/2023.12.03.569481

**Authors:** Jeroen Jansen, Katherine E Moseby, Sebastien Comte, Abbey T Dean, Geoff Axford, David E Peacock, Robert Brandle, Menna E Jones

## Abstract

Re-introduced native animals face major challenges to re-establish themselves in their previous range of distribution when invasive alien predators are present. We study the interactions between a native and an invasive mesopredator across an ecological gradient in the Ikara-Flinders Ranges National Park in South Australia. We placed VHF/ GPS-collars on feral cats (*Felis catus*) and re-introduced western quolls (*Dasyurus geoffroyi*) and analysed their movement patterns, both utilisation distributions and behavioural states, and habitat selection. Despite being able to move anywhere in this open landscape, there is a clear interspecific difference between the two species in their use and preference for landscape elements. Cats prefer grassland and occupy flat areas where rabbit densities are high. Quolls prefer forests and avoid areas of high rabbit and therefore areas of high cat density. The almost complete spatial separation of cats and quolls may reflect different habitat requirements, but the historically broad distribution of quolls suggests that cats may limit quolls continuously to a restricted niche in the Flinders Ranges. This raises the possibility of management interventions that could support a further expansion of quoll occupancy.

## Introduction

Resource availability regulates predator-prey systems and plays an important role in limiting animal populations around carrying capacity and species around their biological optima in their favoured habitat (Stevens 2012). Invasive alien species can disrupt these ecosystem structures, forcing native animals out of their established ecological niches through predation and competition for limited resources (Mooney & Cleland 2001). This disturbance can have severe consequences, leading to a decline in native species abundance, and can even push them to population collapse or extinction (Gurevitch & Padilla 2004). The loss of highly interactive species in the food web, like top predators or pollinators, can disrupt the balance of an ecosystem, triggering trophic cascades that may push the ecosystem to the point of collapse (Pringle *et al*. 2019).

Predation and resource competition interact with environmental productivity and drive the interspecific dynamics amongst native and invasive species at all trophic levels, influencing their abundance and apparent habitat preferences (Chase *et al*. 2002; Chesson & Kuang 2008). While intraguild predation may influence the balance between native and invasive species (Henkanaththegedara & Stockwell 2014), the effect of competition can change the niche characteristics of sympatric species on a large scale (Peers, Thornton & Murray 2013). Two species cannot coexist in the same location if competing for the same limited resources (Gause 1934), as interspecific competitive ability will select for the species with a better ability to exploit the resource (Connell 1961; Sale 1974). The ecological interactions between species thus narrow the potential or fundamental niche to a realized niche (Pearman *et al*. 2008). Coexistence cannot occur, unless both species avoid competition, show considerable differences in their ecological niche (Ayala 1970) or the disadvantaged species restricts its niche to the highly profitable areas (Brown & Wilson 1956).

Invasive animals are placing additional pressure on native animals, restricting their ability to persist in suitable habitat and potentially confining them to reduced realised niches in the long-term (Scheele *et al*. 2017). By reducing habitat quality and extent, human activities and land alteration accelerate and exacerbate the impacts of invasive species and further disadvantage native species (Didham *et al*. 2005). This is particularly prevalent during the periods of resource stress that accompany extended droughts, when animals of the semi-arid zone have already retreated into moister and more vegetated refuge areas (Pavey *et al*. 2017).

In Australia, the combined effects of introduced animals such as European rabbits (*Oryctolagus cuniculus*), red foxes (*Vulpes vulpes*) and feral cats (*Felis catus*), along with European land-use practices of continuous and often over-grazing by domestic livestock, has resulted in the extinctions and population declines of small-mammals. This has led to 11% of endemic terrestrial species being extinct and another 21% threatened (Woinarski, Burbidge & Harrison 2015). Conservation programs across Australia aim to slow biodiversity loss and to recover native animal populations over large scales where possible (Pedler *et al*. 2016).

As part of the large-scale restoration program “Bounceback”, the western quoll (*Dasyurus geoffroyi*), a native marsupial mesopredator, was reintroduced into the Ikara-Flinders Ranges National Park (IFRNP) in South Australia from 2014 to 2016 (Moseby *et al*. 2021a), after successful long-term fox-baiting since 1992 eliminated sustained fox presence within the park (Stobo-Wilson *et al*. 2020). The western quoll is a nationally threatened species that has undergone a range decline of more than 95% since European arrival (Woinarski, Burbidge & Harrison 2014). With a former distribution of almost two thirds of the Australian continent across a wide range of habitats and most of the arid and semi-arid zone, quolls should be able to occupy much of the IFRNP. Quolls remain, however, in habitats with complex structure and refuges, and have not expanded into the open grassy plains (Moseby *et al*. 2021a). Similar to northern quolls in the Pilbara, reasons suspected to limit expansion of the quoll population include predation by feral cats and lack of structural complexity of the vegetation (Hernandez-Santin, Goldizen & Fisher 2022).

In the IFRNP, these threats are connected. The open grassy plains in the IFRNP have the highest abundance of rabbits (Jansen *et al*. 2023) and cats (Stobo-Wilson *et al*. 2020), as well as high grazing and browsing pressure from a range of domestic, feral and native herbivores that severely reduces cover of ground vegetation. Structural complexity of the ground vegetation layer may indirectly influence quoll survival by providing hiding places and food sources as well as enabling them to evade detection or to escape from mammalian predators such as cats (McGregor *et al*. 2015). With their early maturity and rapid reproduction, rabbits can reach high population numbers (Myers 1970). In Australia, rabbits serve as a primary food source for feral cats and foxes in the arid zone (Doherty *et al*. 2015). While introduced predator densities are associated with rabbit densities (Holden & Mutze 2002), foxes have been effectively controlled in the IFRNP and therefore the focus of this study is on cats.

Feral cats are highly adaptable predators found across diverse habitats and climate zones on all continents (Long 2003). They have contributed to the extinction of 63 island species and threaten more than 430 vertebrate species globally (Doherty *et al*. 2016). In Australia, they are considered to have contributed to the extinction of 22 endemic animals and threaten an additional 75 species (Woinarski, Burbidge & Harrison 2015). Cats impact native animals through predation, disease transmission and resource competition (Denny & Dickman 2010). Managing invasive alien species such as feral cats at large scale in open landscapes is complex and challenging (Donaldson, Wilson & Maclean 2017). Cats have the ability to cover vast distances in search of suitable habitat (Jansen *et al*. 2021) and reinvasion can occur rapidly after management efforts (Palmas *et al*. 2020).

To restore native fauna across large landscapes requires ecologically based management approaches, harnessing natural levers where a small action can result in a larger effect elsewhere in the system (Hobbs *et al*. 2011). To identify potential management levers requires a deep understanding of the ecological interactions and dynamics among invasive and native species and the environment. To efficiently manage feral cats and reduce their impact on native wildlife in open landscapes, knowledge of the ecological dynamics amongst multiple species, and how cat presence influences the behaviour and habitat choices of native animals is fundamental. Identifying refuge areas and imminent threats to native wildlife allows to tailor protection and management actions, optimising conservation efforts.

We use GPS tracking of feral cats and the reintroduced native mesopredator, western quolls, to identify their habitat preferences in relation to the abundance of rabbits and other biotic and abiotic features. Our case study is within the IFRNP and surrounding areas in semi-arid South Australia in a large, unfenced landscape. Our objective is to provide detailed knowledge to guide the development of effective conservation management strategies to facilitate native species recovery, including western quolls, in open landscapes in the presence of high densities of invasive predators (cats) and prey (rabbits).

## Methods

### Study areas

The study site is in the 938 km2 IFRNP (31.25◦ S, 138.42◦ E) located in the central Flinders Ranges in semi-arid and arid South Australia (Figure 1). Daily average maximum temperatures are between 25.6–34.2 ◦C in the summer months (October to March), and 16.0–25.4 ◦C in winter (June to August). Rainfall is low (300–400 mm), sporadic, and with higher rainfalls in the winter (BOM 2021). The years between 2017 and 2020 were the driest period on record.

**Figure 1:**
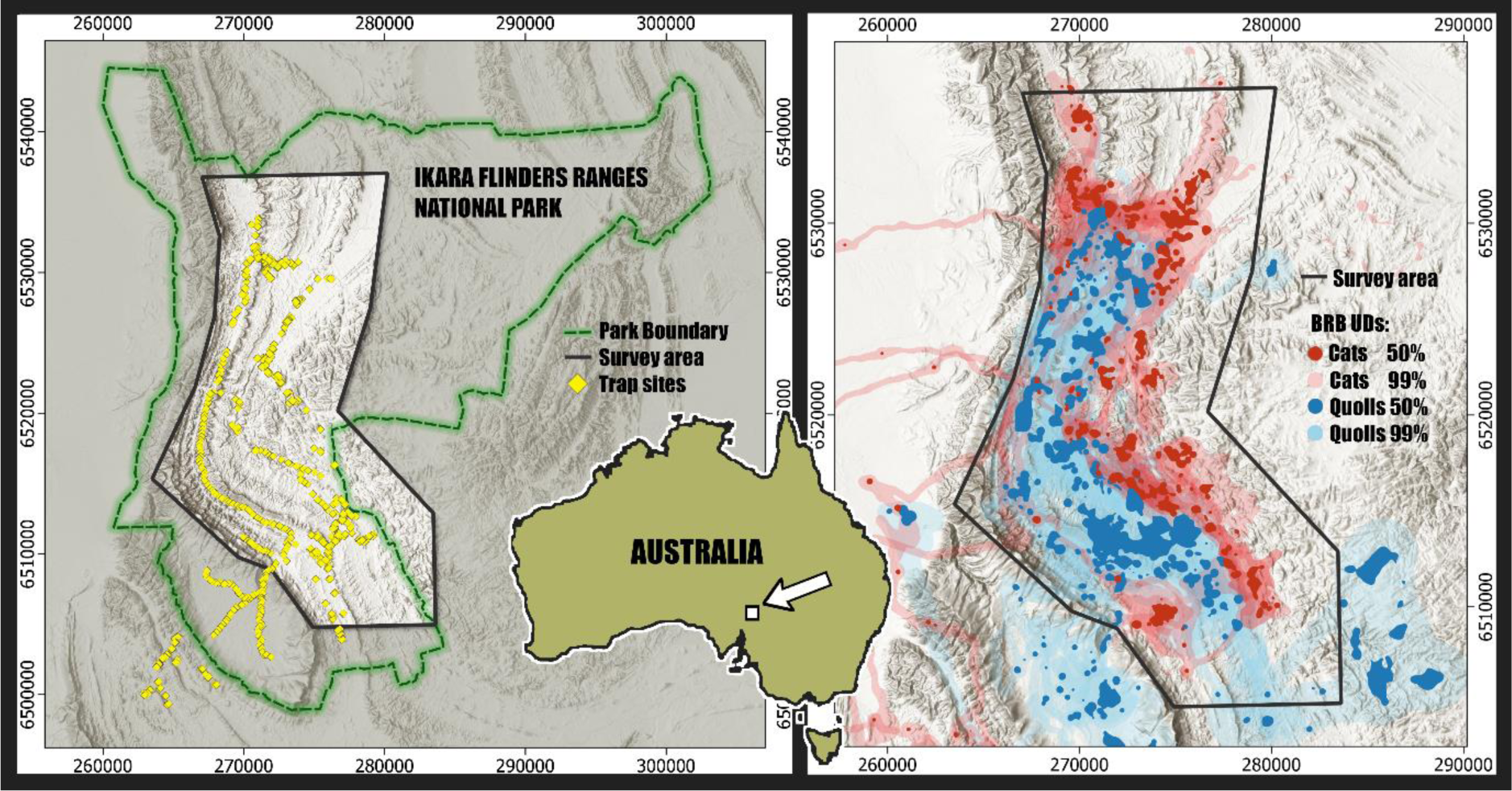
Study site with cage trap locations and utilisation distributions: the left map shows the location of survey area (bright area with black boundary) within the IFRNP (green dashed line) and the trap locations in yellow. The right map shows the biased random bridges utilisation distribution (BRB UD) of western quolls (blue) and feral cats (red), with light shading representing areas of low utilisation (99% BRB) and dark shading high use areas (50% BRB).

### Field data collection

Between 1st April and 10 May in 2019 and between 25 February and 5 May in 2020, we set 40 wire cage traps over about 3500 trap-nights baited with rabbit meat to trap both western quolls and feral cats (Figure 1). Traps were located on animal trails and near creek lines, checked at first light, and closed during the daytime. All trapped animals were weighed and sexed and later released at the point of capture into vegetation cover. If meeting the minimum weight (cats 1000 g, quolls 933 g) and health requirements, captured animals were fitted with GPS collars before release. Feral cats were fitted with Telemetry Solutions 4000ER GPS/VHF collars (weight: 78-81g) or an ATS G10-M1740 collar (50g) programmed to record the location of the animals every 15 minutes (Table 1). Quolls were fitted with Nano GPS/VHF collars (weight: 28-31g) using embroidery thread as a weak link (Morrant *et al*. 2022). Due to their smaller and more limited battery size and life, the collars on quolls were programmed to record the location of the animals every 15 minutes from one hour before sunset (17:00) to one hour after sunrise (7:00). Quolls that were to be fitted with GPS collars were held in a bag within a covered trap in a cool and dark place during daylight and released after sunset on the same day. Recording of GPS data started 2 days after release of the animal to avoid recording potential effects of capture on the behaviour of the animal.

**Table 1:**
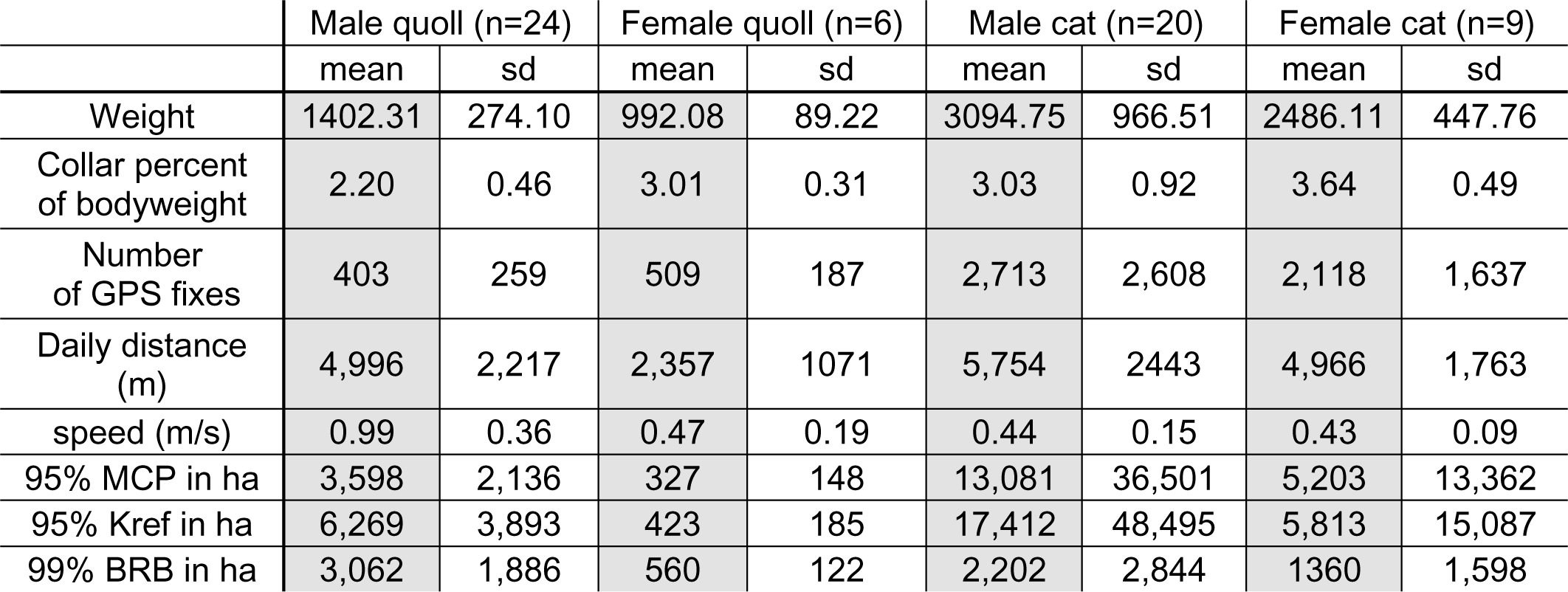
Metrics of the movement data from GPS-collared cats and quolls in the IFRNP in South Australia.

Collared animals were located daily using a car-mounted VHF omni-antenna and a hand-held 5 element 150 MHz folding antenna (both Advanced Telemetry Systems). We additionally conducted weekly monitoring flights in a Cessna 172 plane with a Civil Aviation Safety Authority-approved VHF antenna (Faunatech) affixed to the wing strut. Due to COVID-19 restrictions in 2020, the use of aircraft for tracking only started in mid-May. Tracking flight paths followed parallel lines across and extending beyond the entire study area, to find animals that wandered beyond the park boundaries (Jansen *et al*. 2021). Once a month, GPS fixes stored on board the collars were remotely downloaded using a Bluetooth base station.

All collared cats were euthanised by qualified personnel at the end of the study (four months after the date of the first collar deployment) in accordance with the scientific permit requirements. With most of the quoll collars, the weak link (embroidery thread) broke, and the collars fell off before the end of study. To ensure all collars were removed, quolls were re-trapped at the end of study, weighed, checked for potential abrasions or wounds resulting from wearing the collar, and the collar removed if it had not already fallen off.

### Data handling

Animal location fixes obtained from the GPS collars were cleaned following Bjorneraas *et al*. (2010). We excluded data sets that had less than 10 days of recorded data and only kept location fixes recorded as being in three dimensions by the collars and we filtered out location fixes obtained with less than three satellites or with a horizontal dilution of precision (hdop) above seven. We then screened the remaining tracking data for outlier GPS points that resulted in speeds higher than 20km/h over the 15min step. We finally removed all locations where successive fixes indicated the collar was stationary, which corresponded to when an animal was trapped, following the activation of the drop-off mechanism on the collar or the death of the animal. We used these stationary above-ground locations to estimate the precision of the GPS collars by calculating the deviation from the centroid of the points. We used a random walk model (McClintock & Michelot 2018) to generate up to two virtual locations of the animals when true GPS locations were missing. Any track section of less than four continuous GPS points was discarded.

Six environmental parameters were derived from publicly available gridded layers (i.e. raster) at 1-second resolution (approximately 25×30m) or from publicly available spatial polygon data. All environmental data (raster and polygons) were converted to UTM and resampled to a unique 25×25m grid across the whole study area. We included the following layers. A) Geology (SARIG 2021) was combined into six general geological categories: quartzite (1), sandstone (2), deposits (3), siltstone (4), limestone (5) and shale (6). B) The vegetation cover (VC) was reduced from 16 originally available classes to 5 categories (Willoughby *et al*. 2018): disturbed ground (1), nonwoody vegetation/ grassland/ low shrubs (2), sparse vegetation (3), low native vegetation/ samphire species (4), and forests/ woodlands (5). C) The normalised difference vegetation index (NDVI) was calculated from Sentinel 2 satellite data (U.S. Geological Survey 2022) as a continuous variable ranging from 0 to 1 using the near infra-red (NIR) and red bandwidth, and the following formula: NDVI = (NIR - Red) / (NIR + Red). D) Terrain Ruggedness Index (TRI) was calculated using the raster package (Hijmans *et al*. 2015) from a hydrologically enforced digital elevation model (Wilson *et al*. 2011). E) The Prescott Index (PI) uses long-term average precipitation and evaporation potential to estimate the water balance in different parts of the landscape (Gallant & Austin 2012a). F) The Topographic Wetness Index (TWI) estimates the relative wetness of parts of the landscape by the size of catchment and the slope (Gallant & Austin 2012b). In addition to these six environmental parameters, we used maps of the potential rabbit abundance derived from previous ground surveys across the study area (see Jansen et al. (2023) for the detailed methods). For each GPS location of cats and quolls, we extracted the values of the six environmental parameters and potential rabbit abundance for the cell (25×25m) containing the GPS location. We checked for collinearity of the environmental variables with a Spearman test.

### Statistical analysis

Data analyses were done using *R* version 4.0.3 (R Core Team 2020), *R studio* version 1.3.1093 (RStudio Team 2020) and *QGIS* version 3.6.0 (QGIS Development Team 2020).

For each collared individual, we first calculated the linear distances (step length) between points that were consecutively recorded without gap. For each step, we then calculated the speed of the animal (m/s) as the step length (m) divided by the step duration (seconds). The daily distance moved as the sum of step length between noon and noon the next day (24 hours) were also calculated. For each species, generalised linear mixed models (GLMM) were used to describe the influence of sex, bodyweight, and the time of the year, parameterised as the day of the year on the daily distance moved, with individual as a random effect. All models were fitted in *glmmTMB* (Magnusson *et al*. 2017).

Home ranges were calculated for each collared animal as biased random bridges (BRB). The BRB calculation is a movement-based method that considers the activity time between successive locations of serially correlated relocations of an animal to estimate utilisation distributions (UDs), and thus improves the biological relevance of UD estimates (Benhamou 2011). For comparison with older publications, we calculated the 95% minimum convex polygon (MCP) and 95% Kernel density home ranges (Kref) with reference bandwidth (default; href). All home ranges were calculated using the *adehabitat* package version 0.4.19 (Calenge 2006). We compared the sizes of home ranges between the different species and sexes using a t-test and calculated the area of overlap between individual home ranges using the 99%BRB home range polygons. As there were insufficient GPS data available, we did not test for an effect of year.

Habitat selection ratios were calculated for each collared individual using the ratio of available to used habitat (Aebischer, Robertson & Kenward 1993). Available habitat was determined by creating a buffer of 2000m around the individual contour of the 99% BRB home range. This buffer distance is less than 90% of the lowest mean daily average distance (2257m/ female quoll) of all collared animals during this study (Table 1). All animals should therefore be able to walk this distance in a day. The sum of the grid cells for each environmental parameter contained in this area was used as the available habitat. We compared the distribution of these environmental variables with the sum of grid cells of each variable present within the 50% UD for each individual quoll or cat. The habitat selection ratio was then calculated as used divided by the available habitat for each environmental variable. The resulting individual values were then combined for each sex of the two species.

To compare the environmental influences on the movement behaviour of each species and sex, we used hidden Markov models (HMM), which analyses the movement angles and step lengths between relocations, to categorise movement behaviour into states and calculate the stationary probability of each state. The *momentuHMM* package (McClintock & Michelot 2018) was used to determine first whether a 2-state or 3-state model best represented the observed distributions of different movements. The relative influence of the environmental parameters was then tested with a candidate model set including different parameters. We used multi-model inference and Akaike’s information criterion, following the model selection process in Richards (2008) to select the most parsimonious model. The Viterbi algorithm on the fitted model was used to determine the most likely state sequence.

## Results

A total of 17 cats and 11 quolls in 2019, and 19 cats and 24 quolls in 2020, were of sufficient minimum weight to be fitted with GPS collars (Table 1). Cats carried the collars for between 10 and 128 days (Supplementary Table 1 (S1)), and quolls for between 14 and 71 days (S 2). After GPS data cleaning, there were between 104 and 8056 GPS points for individual cats (S 3), and between 53 and 891 GPS points for quolls (S 4). The mean precision of the GPS/VHF collars was 13.5 m ± 7.8 for cats (S 5) and 20.4 m ± 5.1 for quolls (S 6).

There was no effect of body weight on the daily distance moved by male and female quolls but a strong influence of sex: males move more than females (Figure 2, S 7). Additionally, the daily distance moved had a non-linear response to time with a different effect for male and female quolls. Male quolls increased the daily distance they moved between the beginning of the survey period in April and remained moving further distances after the end of May, whereas females decreased their distances until mid-May and increased the distance thereafter. For cats, a weak effect of weight on the movement distance is detected, where heavier animals move slightly further distances, but the day of the year and the sex did not influence the daily distance moved (S8).

**Figure 2:**
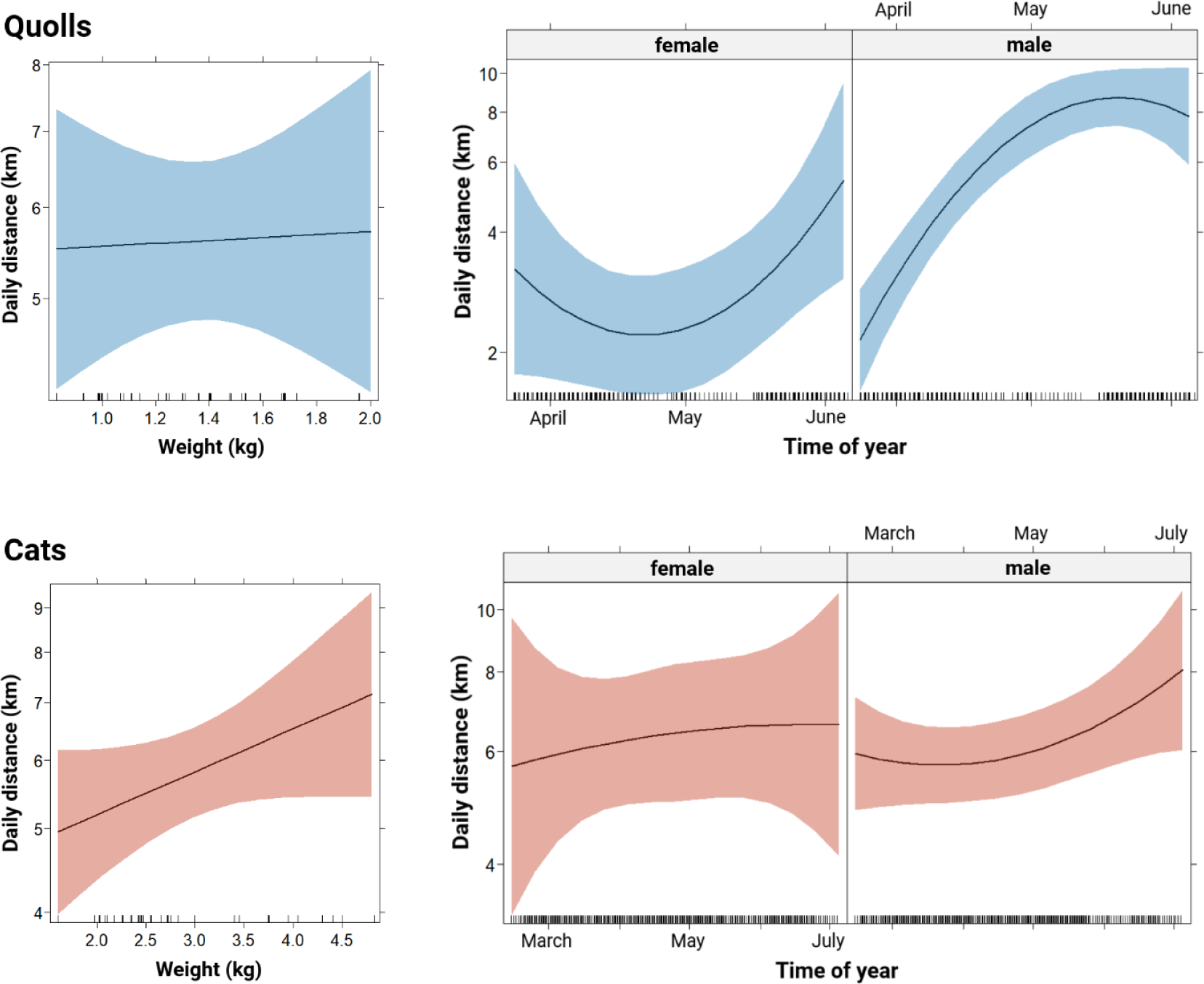
Movement data of feral cats and western quolls within the Ikara-Flinders Ranges National Park in South Australia. The left-hand graphs show the median distance moved per day (sum of step lengths) for each species in relation to bodyweight. The right-hand graphs show the distance moved per day across the survey period for each sex and species. Confidence intervals are displayed in colour.

Home ranges were between 392 ha and 705 ha (mean 560 ha) for female quolls and between 744 ha and 7,217 ha (mean 3,062 ha) for male quolls (S4), and between 545 ha and 5,588 ha (mean 1,598 ha) for female cats and between 51 ha and 12,533 ha (mean 2,844 ha) for the male cats (S3). Home ranges (p = 4.21e-07) and travel speeds (p = 1.47 e-04) differ between male and female quolls, whereas there is no difference in these factors between male and female cats (S9).

The home ranges of cats and quolls are spatially separated (Figure 1). Of the individual quoll home ranges, 95.6% of the 50% BRB home ranges do not overlap with a home range of a cat and just 0.5% show an overlap of more than 5% of the home range area (S 10). There is more overlap between home ranges of pairs of individual quolls in the 50% BRB and 99% BRB than between individual cats, or cats and quolls (S 10 and 11).

In this environment, quolls and cats show differences in their habitat selection ratios in most of the environmental factors analysed (Figure 3). They select for contrasting densities of rabbit abundance, distinctive vegetation cover (disturbed ground vs forest) and prefer different geology layers. Quolls strongly select against medium to high rabbit densities, grasslands and limestone and shale geologies. They prefer low rabbit densities, forested areas, and siltstone layers. Additionally, female quolls differ from cats and male quolls in their preference for low to high terrain ruggedness and topographic wetness index. Cats show a strong selection for medium to high rabbit densities, grasslands and limestone and shale layers. They prefer not to be in forests, siltstone layers and areas with low rabbit densities.

**Figure 3:**
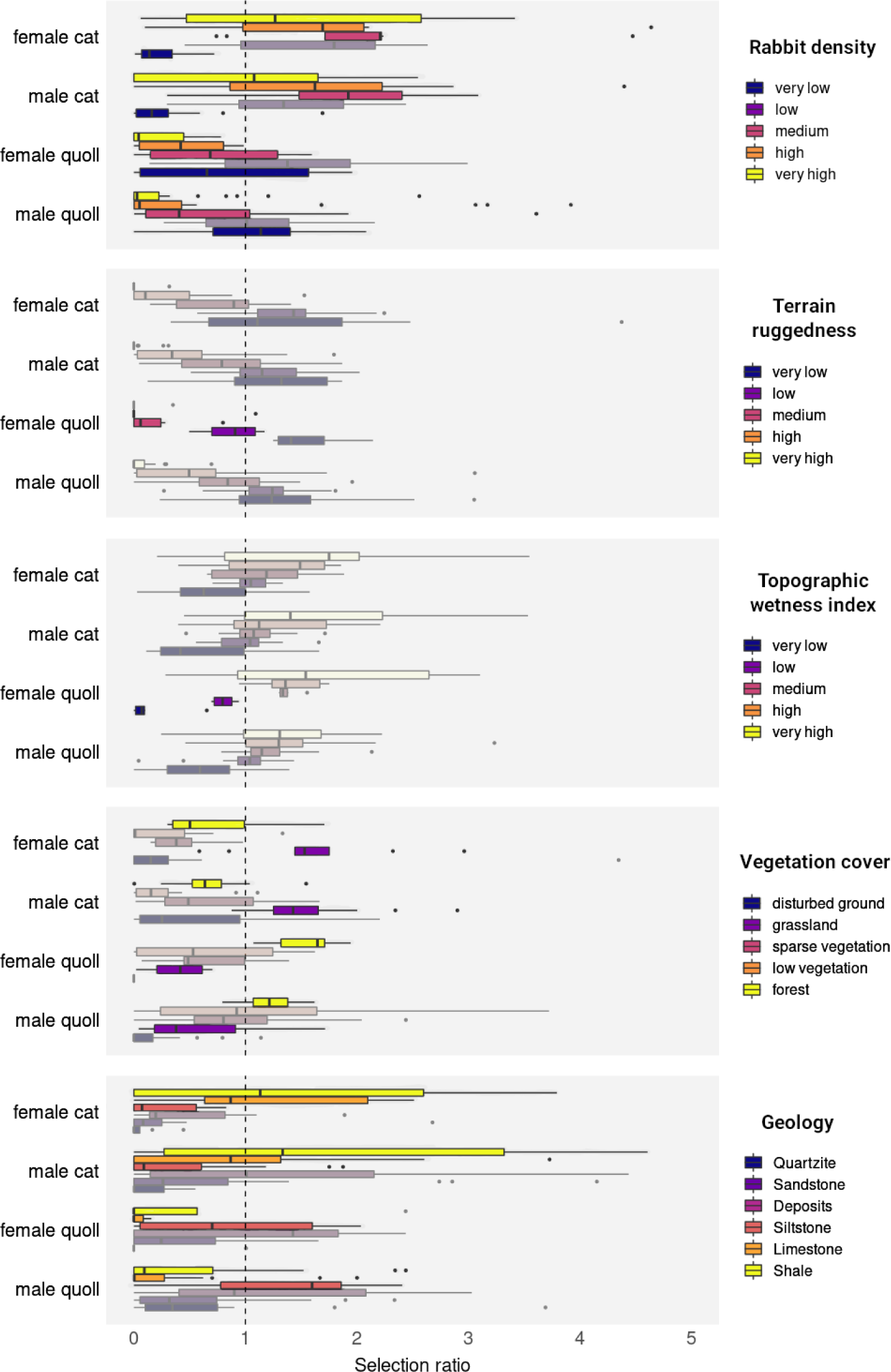
Habitat selection ratios of feral cats and western quolls for five different environmental parameters within the Ikara-Flinders Ranges National Park in South Australia. Sexes are analysed and plotted separately. Outliers above 5 have been removed for clearer visual representation; the full graph is in the supplementary material. Values are calculated using UD values lower than 0.5 and the available habitat within the buffer around the individual home range. The dotted line represents no selection, values greater than are selected for, and values less than are selected against. Where there are species differences, these are highlighted using strong colours for easier visualisation.

The best representation of the observed distributions of different movements was a 3-state model for the quolls, and a 2-state model for the cats (Figure 4; S 13 and S14). The three states of quolls can be differentiated into transiting (fast linear movement), foraging (local tortuous movement) and stationary. With the cats, a further separation of the behaviours to more than 2 states cannot be done and are therefore described as moving (displacement happening) and stationary. For each species and sex, different environmental factors influence their movement decisions (Figure 4). The most parsimonious models describing movement states included the Prescott Index and rabbit abundance for female quolls (S 15), the terrain ruggedness index for male quolls (S 16), geology and the terrain ruggedness index for female cats (S 17), and the Prescott Index and geology for male cats (S 18).

**Figure 4:**
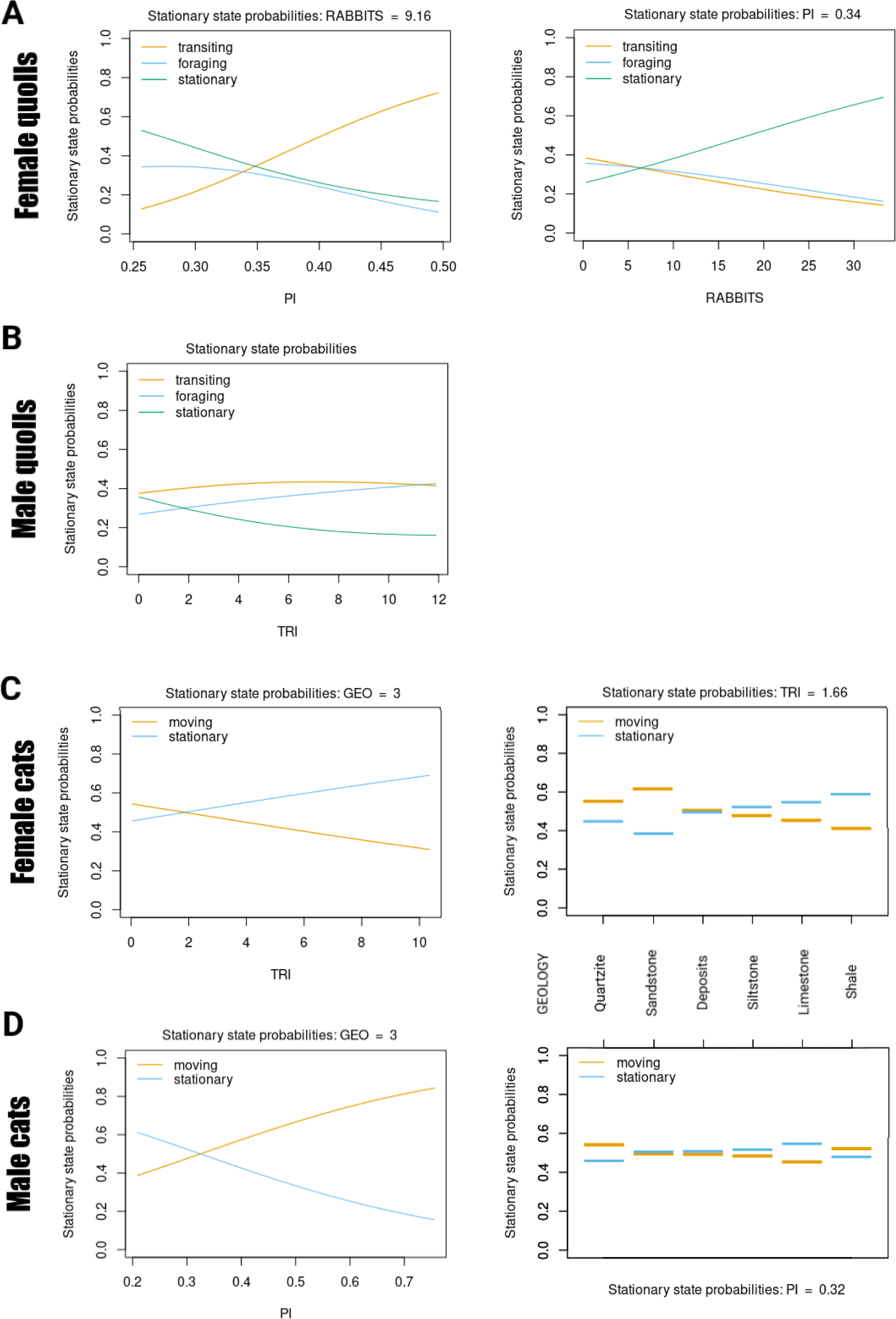
Stationary state probabilities of each movement state across environmental parameters for feral cats and western quolls in the Ikara-Flinders Ranges National Park in South Australia. Each graph shows an environmental parameter from the top models describing the movement state. Females and males of each species are presented separately. PI = Prescott Index, Rabbits = rabbit abundance, TRI = terrain ruggedness index and geology.

Female quolls increased their transiting movements and decreased stationary and foraging behaviours with increasing Prescott Index, or higher moisture (Figure 4a). They also decreased their transiting and foraging movements with increasing rabbit density, they increased stationary behaviour at higher rabbit densities which means when present in these parts of the landscape they reduced movement. As terrain ruggedness increases, the probability of male quolls remaining stationary decreased, the probability of them switching to a transiting movement pattern stayed at a similar level and the probability of switching to a foraging pattern slightly increased (Figure 4b). Female cats were more likely to remain stationary as ruggedness of terrain increases and when on in siltstone, limestone and shale geology, while they were more likely to be moving where ruggedness is low and when in quartzite and sandstone (Figure 4c). Male cats were more likely to move with increasing Prescott Index and when on quartzite and shale geologies, whereas they were more likely to remain stationary with low Prescott Index and when on siltstone and limestone (Figure 4d).

## Discussion

The movement data from the GPS-collared quolls and cats show clear interspecific differences in space use across the survey area. Even though some of the traps captured both species, the overlap in use of space after release of the animals is minimal. Individuals of both quolls and cats did exploratory movements into areas mainly occupied by the other species, but they never remained there for longer than a few hours. The habitat selection ratios strengthen this image of an interspecific divide with clear differences between the species in their preferences for certain environmental variables such as the vegetation cover, geology, and rabbit densities. All the animals collared show broadly similar movement behaviours, with comparable sized home ranges and distances per day travelled, despite that the cats that travelled long distances (Jansen *et al*. 2021) were included in the data set. The large differences in daily distances travelled between male and female quoll, along with male quolls having the largest home ranges of any species/sex combination of the carnivores collared, might be attributed to the study period in both years coinciding with the mating season (Stead-Richardson *et al*. 2001).

With their quite different preferences for distinctive environmental variables, there are several alternative hypotheses for the patterns: 1) distinct realised niches as outlined above; 2) predatory or competitive exclusion of quolls by cats shaping a current reduced realised niche; 3) the areas of current quoll occupancy providing a temporal refuge in a droughted and over-grazed landscape; and 4) the potential for a long-term reduced niche driven by ongoing over-grazing and high rabbit and cat densities. Cats and foxes are both regularly baited in the IFRNP, which suppresses cat populations and effectively eliminates foxes, and the quolls have been reintroduced relatively recently, with their populations still building (Moseby *et al*. 2021a).

The sharp demarcation between the areas occupied by cats and quolls in the IFRNP and the habitat selection ratios indicates that cats may limit the area and habitats that could potentially be occupied by quolls. Cat distribution and abundance in this study and more generally across Australia is positively associated with rabbit abundance, their preferred food resource (Molsher, Newsome & Dickman 1999; Doherty *et al*. 2015). In the harsh climate of hot semi-arid Australia, seasonally high physiological stresses will exert strong pressure for predators to forage optimally, on prey with higher energy returns (Schoener 1971), and to linger around these resources (Shelford 1931). In arid environments, feral cats are biophysically constrained and must seek refuge in periods of high temperatures (Briscoe *et al*. 2022). This suggests that foraging activity in such environments must be efficient and the association with areas of high rabbit density could be due to foraging efficiency and/or the presence of rabbit burrows that provide a thermal refuge. With intensification of cat control in the IFRNP leading to a reduced number of cats, space use is now tightly focussed on the areas of highest rabbit density. The areas that are sub-optimal for cats, partly because of their low rabbit densities, are occupied by the increasing quoll population.

It is possible that cats exclude quolls through predation pressure. Cats have previously been identified as a threat to the quolls in the IFRNP (Moseby *et al*. 2021b). They can adjust their diel activity pattern to the activity time of quolls at the time of year when juveniles become independent, creating a ‘predator pit’ by killing almost the entire annual cohort of young quolls (Fancourt *et al*. 2015). The ongoing expansion of quolls in the Flinders Ranges, though, indicates that they can sustain the current predation pressure from cats in the areas they are occupying (Rob Brandle pers comm.). Adult quolls are probably better able to avoid predation from cats than juveniles, as suggested by experimental study of risk perception in the similar eastern quoll (*D. viverrinus*) (Jones, Smith & Jones 2004). Therefore, predation pressure alone may not explain the spatial separation of the species entirely.

There are no indications that quolls in the IFRNP are in direct competition with cats for food or habitat resources. Both quolls and cats prey on rabbits, but quolls appear to be very good at finding rabbits when they are at low density, whereas cats appear to require high concentrations of rabbits (Moseby, Jensen & Tatler 2022). Western quolls originally occupied a diverse range of forest, shrubland and desert habitats, although their last wild distribution in Western Australia, which may be a reduced realised niche limited by cats and foxes, is in forest (Serena & Soderquist 1989). Western quolls in the IFRNP should be able to use the open grassland habitat currently dominated by cats and rabbits. Western quolls successfully breed and establish themselves in open habitat when introduced predators are excluded (West, Tilley & Moseby 2020), and eastern quolls prefer open grassland habitats in Tasmania (Jones & Barmuta 2000). Yet, we did not record quolls using or attempting to establish themselves in the areas used by cats. They avoided these areas of open grasslands and high rabbit-disturbance, restricting themselves to areas with higher moisture content and greater vegetation density and complexity (Brandle 2001). In the arid zone, areas of higher plant diversity and vegetation complexity should support high invertebrate diversity and abundance (Crisp, Dickinson & Gibbs 1998; Knops *et al*. 1999), fluctuating with rainfall and vegetation cover (Kwok *et al*. 2016). These denser vegetated areas seem to be optimal quoll habitat perhaps because they provide sufficient dietary requirements.

Another possible explanation is that the areas occupied by quolls during the study represent a temporal refuge (Pavey *et al*. 2017), even without needing to consider predation risk from cats. Quoll activity is currently focussed on areas that provide food and refuge, avoiding predation by cats and open grasslands where vegetative cover has been denuded, exacerbated by the exceptional drought during which this study was conducted. The formerly broad distribution of western quolls across the Australian arid and semi-arid zone, indicates they are physiologically well-adapted to arid locations (Morton 1982), likely more so than feral cats. Whereas cats are reliant on rabbits, which as a food source have a high water content, the smaller quolls consume a wide variety of prey, including a high proportion of invertebrates, in both Western Australia and in the IFRNP (Soderquist & Serena 1994; Moseby, Jensen & Tatler 2022). These dietary differences allow quolls to occupy areas that would be not ideal for cats. If these factors are the predominant driver of quoll occupancy, and quolls can sustain predation pressure from cats, then quolls should be able to expand into open grassland areas once the drought breaks. This recovery would take time because quolls breed only once a year and are not capable of the irruptions or rapid relocation of populations to surrounding areas that typify the species in Pavey *et al*. (2017) on which the refuge concept was developed. A complication for quolls to reoccupy open grassland is the rapid increase in rabbit numbers following rainfall and vegetation growth. Without rabbit control, denudation of vegetation structure in open grasslands will be ongoing and cat densities will increase.

The current realised niche of quoll occupancy in the IFRNP, which may represent a refuge distribution, could become a long-term reduced niche (Scheele *et al*. 2017) if the current rabbit densities persist, even in times of good rainfall. Rabbit populations respond positively to rainfall preventing the recovery of vegetation and supporting high cat densities. While quolls may be able to survive low to intermediate cat densities, they may be limited in the long term by high predation pressure and limited food in the vast areas of open grasslands. Cats in the IFRNP are found in all areas of the park and, prior to intensification of cat control, were found at higher densities in areas now used by the quolls (Moseby *et al*. 2021a). Our results on the associations between cat space use and rabbit abundance and the history of cat control in the IFRNP suggest this situation is dynamic. This means it may be possible to shift the distribution and abundance of invasive predators in the IFRNP by applying management actions focussed on reducing the abundance of rabbits, thereby altering the ecological dynamics to favour native animals (Dean *et al*. 2023).

### Future research and management

The next steps are to build on the knowledge generated in this study to test the hypotheses of the ecological drivers of quoll distribution and abundance and identify and design ecologically based management levers with the aim to provide options and scenarios for conservation managers. These management approaches need to provide the conditions for quolls to maintain self-sustaining populations and to increase the rate at which they can expand their area of occupancy out from the safe refuges into the broader landscape of the Flinders Ranges, including crossing gaps of open grassland between isolated ranges and even occupying open grasslands. This could be done by reducing the rabbit abundance, an important resource driving cat density (Stobo-Wilson *et al*. 2020). Reducing rabbits will allow native vegetation to recover, providing structural complexity that native animals require to avoid predation and bring cat numbers down by reducing their capacity for population build up (Courchamp, Langlais & Sugihara 2000). Previous research (McGregor *et al*. 2019) and our findings indicate that in the semi-arid zone, controlling rabbits in one area will likely make cats move to adjacent areas with high rabbit abundance, rather than ongoing prey-switching.

The success of the re-establishment of the western quoll in this area is based on the current negligible fox and reduced cat density. Even when rabbit control can be implemented as an ecological lever to reduce cat activity and enhance and connect habitat for quolls, the lethal management pressure on foxes and possible also cats needs to be maintained.

## Conclusions

We demonstrate with this case study that the space use and habitat preferences of a successfully reintroduced native predator (western quoll) may be defined as a consequence of elevated numbers of an invasive prey species (rabbit) supporting high densities of an invasive predator (cat), a situation potentially exacerbated by drought and over-grazing. Changes in the occupancy of both cats and quolls indicate that this system is dynamic. This lends this semi-arid system to management manipulations grounded in food web interactions to expand and connect habitat for the native predator. Using the mechanisms of this invasive predator prey dynamic, methods to tilt the balance towards better native animal protection can be achieved.

Invasive predators are often co-introduced with invasive prey across the world, leading to spill-over or hyperpredation on native species. Finding ecological levers to nudge the system towards facilitating native species and disadvantaging invasive species is cost-efficient and an effective way of achieving conservation goals. Ecologically based conservation management can be accomplished with a detailed understanding of the habitat preferences and movement dynamics amongst the native and invasive predator and prey species in the system. This is especially relevant in vast open landscapes where occupancy and abundance levels are dynamic and not contained by artificial or natural boundaries but equally as important on a small scale during local conservation efforts or for small populations.

## Author Contributions

Conceptualization and methodology: JJ, SC, KEM, RB, and MEJ; formal analysis: JJ, SC; data curation: JJ, ATD, GA, DEP, KEM, and RB; resources: JJ, GA, RB and MEJ; writing - original draft preparation: JJ, SC and MEJ; writing - review and editing: all authors; visualization: JJ; funding acquisition: GA, RB and MEJ; All authors have read and approved the published version of the manuscript.

## Acknowledgements

We thank Luke Yates, Ben Halliwell, Leon Barmuta, Cyril Scomparin and Rahil Amin for help during analysis; Chris Johnson for suggestions on study design; Cat Lynch and volunteers for assistance during fieldwork; Catherine Bresnehan and the SNS administration office for tremendous help with paperwork, Iain Dunk and the Department for Environment and Water for providing data; Adnyamathana members of the IFRNP co-management board for permission to do fieldwork on their country, and National Parks and Wildlife Service for ongoing support and access to park facilities. This research was supported by use of the NeCTAR Research Cloud, a collaborative Australian research platform supported by the NCRIS-funded Australian Research Data Commons (ARCR 2022).

This study is a collaboration between the University of Tasmania (UTAS) and the Department of Environment and Water SA (DEW) with the primary Bounceback research project funded by an Australian Research Council Discovery scheme grant to MEJ (DP170101653), and funding support from the Foundation for Australia’s Most Endangered Species (FAME) for DEW to reintroduce western quolls and brush-tailed possums into IFRNP. Jeroen Jansen was supported by a Tasmania Graduate Research and a Tuition Fee Scholarship. Research in the IFRNP was authorised under SA scientific research permit Q26950-1 and UTAS animal ethics permit A18535.

## Data Availability Statement

The data presented in this study will be available on request from the corresponding author.

## References

Aebischer, N.J., Robertson, P.A. & Kenward, R.E. (1993) Compositional Analysis of Habitat Use From Animal Radio-Tracking Data. Ecology, 74, 1313–1325.

ARCR (2022) NeCTAR Research Cloud. ardc.edu.au.

Ayala, F.J. (1970) Competition, Coexistence, and Evolution. Essays in Evolution and Genetics in Honor of Theodosius Dobzhansky (eds M.K. Hecht & W.C. Steere), pp. 121–158. Springer US, Boston, MA.

Benhamou, S. (2011) Dynamic approach to space and habitat use based on biased random bridges. PLoS One, 6, e14592.

Bjorneraas, K., Van Moorter, B., Rolandsen, C.M. & Herfindal, I. (2010) Screening Global Positioning System Location Data for Errors Using Animal Movement Characteristics. The Journal of Wildlife Management, 74, 1361–1366.

BOM (2021) Australian Government Bureau of Meteorology: Climate Statistics for Australian Locations.

Brandle, R. (2001) A biological survey of the Flinders Ranges South Australia. Department for Environment and Heritage, Adelaide.

Briscoe, N.J., McGregor, H., Roshier, D., Carter, A., Wintle, B.A. & Kearney, M.R. (2022) Too hot to hunt: Mechanistic predictions of thermal refuge from cat predation risk. Conservation Letters, 15, e12906.

Brown, W.L. & Wilson, E.O. (1956) Character Displacement. Systematic Zoology, 5, 49–64.

Calenge, C. (2006) The package “adehabitat” for the R software: A tool for the analysis of space and habitat use by animals. Ecological Modelling, 197, 516–519.

Chase, J.M., Abrams, P.A., Grover, J.P., Diehl, S., Chesson, P., Holt, R.D., Richards, S.A., Nisbet, R.M. & Case, T.J. (2002) The interaction between predation and competition: a review and synthesis. Ecology letters, 5, 302–315.

Chesson, P. & Kuang, J.J. (2008) The interaction between predation and competition. Nature, 456, 235–238.

Connell, J.H. (1961) The influence of interspecific competition and other factors on the distribution of the barnacle Chthamalus stellatus. Ecology, 710–723.

Courchamp, F., Langlais, M. & Sugihara, G. (2000) Rabbits killing birds: modelling the hyperpredation process. Journal of Animal Ecology, 69, 154–164.

Crisp, P.N., Dickinson, K.J.M. & Gibbs, G.W. (1998) Does native invertebrate diversity reflect native plant diversity? A case study from New Zealand and implications for conservation. Biological Conservation, 83, 209–220.

Dean, A.T., Brandle, R., Barmuta, L.A., Jones, M.E. & Jansen, J. (2023) Rabbit warrens: an important resource for invasive alien species in semi-arid Australia. Wildlife Research, -.

Denny, E.A. & Dickman, C. (2010) Review of cat ecology and management strategies in Australia. Invasive Animals Cooperative Research Centre, Canberra.

Didham, R.K., Tylianakis, J.M., Hutchison, M.A., Ewers, R.M. & Gemmell, N.J. (2005) Are invasive species the drivers of ecological change? Trends Ecol Evol, 20, 470–474.

Doherty, T.S., Davis, R.A., van Etten, E.J.B., Algar, D., Collier, N., Dickman, C.R., Edwards, G., Masters, P., Palmer, R. & Robinson, S. (2015) A continental-scale analysis of feral cat diet in Australia. Journal of Biogeography, 42, 964–975.

Doherty, T.S., Glen, A.S., Nimmo, D.G., Ritchie, E.G. & Dickman, C.R. (2016) Invasive predators and global biodiversity loss. Proc Natl Acad Sci U S A, 113, 11261–11265.

Donaldson, L., Wilson, R.J. & Maclean, I.M.D. (2017) Old concepts, new challenges: adapting landscape-scale conservation to the twenty-first century. Biodivers Conserv, 26, 527–552.

Fancourt, B.A., Hawkins, C.E., Cameron, E.Z., Jones, M.E. & Nicol, S.C. (2015) Devil declines and catastrophic cascades: is mesopredator release of feral cats inhibiting recovery of the eastern quoll? PLoS One, 10, e0119303.

Gallant, J. & Austin, J. (2012a) Prescott Index Derived from 1 ″SRTM DEM-s. V2, CSIRO. Data Collection.

Gallant, J. & Austin, J. (2012b) Topographic Wetness Index derived from 1” SRTM DEM-H. V2. 11 CSIRO. Data Collection.

Gause, G.F. (1934) The struggle for existence. Williams and Wilkins Baltimore, Maryland.

Gurevitch, J. & Padilla, D.K. (2004) Are invasive species a major cause of extinctions? Trends Ecol Evol, 19, 470–474.

Henkanaththegedara, S.M. & Stockwell, C.A. (2014) Intraguild predation may facilitate coexistence of native and non-native fish. Journal of Applied Ecology, 51, 1057–1065.

Hernandez-Santin, L., Goldizen, A.W. & Fisher, D.O. (2022) Northern quolls in the Pilbara persist in high-quality habitat, despite a decline trajectory consistent with range eclipse by feral cats. Conservation Science and Practice, 4, e12733.

Hijmans, R.J., Van Etten, J., Cheng, J., Mattiuzzi, M., Sumner, M., Greenberg, J.A., Lamigueiro, O.P., Bevan, A., Racine, E.B. & Shortridge, A. (2015) Package ‘raster’. R package, 734, 473.

Hobbs, R.J., Hallett, L.M., Ehrlich, P.R. & Mooney, H.A. (2011) Intervention Ecology: Applying Ecological Science in the Twenty-first Century. BioScience, 61, 442–450.

Holden, C. & Mutze, G. (2002) Impact of rabbit haemorrhagic disease on introduced predators in the Flinders Ranges, South Australia. Wildlife Research, 29, 615–626.

Jansen, J., Jansen, J., Dean, A.T., Brandle, R., Peacock, D.E. & Jones, M.E. (2023) High-resolution mapping of rabbit (Oryctolagus cuniculus) densities for targeted conservation management. Journal of Applied Ecology, **n/a**.

Jansen, J., McGregor, H., Axford, G., Dean, A.T., Comte, S., Johnson, C.N., Moseby, K.E., Brandle, R., Peacock, D.E. & Jones, M.E. (2021) Long-Distance Movements of Feral Cats in Semi-Arid South Australia and Implications for Conservation Management. Animals (Basel), 11, 3125.

Jones, M.E. & Barmuta, L.A. (2000) Niche differentiation among sympatric Australian dasyurid carnivores. Journal of Mammalogy, 81, 434–447.

Jones, M.E., Smith, G.C. & Jones, S.M. (2004) Is anti-predator behaviour in Tasmanian eastern quolls (Dasyurus viverrinus) effective against introduced predators? Animal Conservation, 7, 155–160.

Knops, J.M.H., Tilman, D., Haddad, N.M., Naeem, S., Mitchell, C.E., Haarstad, J., Ritchie, M.E., Howe, K.M., Reich, P.B., Siemann, E. & Groth, J. (1999) Effects of plant species richness on invasion dynamics, disease outbreaks, insect abundances and diversity. Ecol Lett, 2, 286–293.

Kwok, A.B.C., Wardle, G.M., Greenville, A.C. & Dickman, C.R. (2016) Long-term patterns of invertebrate abundance and relationships to environmental factors in arid Australia. Austral Ecology, 41, 480–491.

Long, J.L. (2003) Introduced mammals of the world: their history, distribution, and influence. CSIRO publishing.

Magnusson, A., Skaug, H., Nielsen, A., Berg, C., Kristensen, K., Maechler, M., van Bentham, K., Bolker, B., Brooks, M. & Brooks, M.M. (2017) Package ‘glmmtmb’. R Package Version 0.2. 0, 25.

McClintock, B.T. & Michelot, T. (2018) momentuHMM: R package for generalized hidden Markov models of animal movement. Methods in Ecology and Evolution, 9, 1518–1530.

McGregor, H., Legge, S., Jones, M.E. & Johnson, C.N. (2015) Feral Cats Are Better Killers in Open Habitats, Revealed by Animal-Borne Video. PLoS One, 10, e0133915.

McGregor, H., Moseby, K., Johnson, C.N. & Legge, S. (2019) The short-term response of feral cats to rabbit population decline: Are alternative native prey more at risk? Biological Invasions, 22, 799–811.

Molsher, R., Newsome, A. & Dickman, C. (1999) Feeding ecology and population dynamics of the feral cat (Felis catus) in relation to the availability of prey in central-eastern New South Wales. Wildlife Research, 26, 593–607.

Mooney, H.A. & Cleland, E.E. (2001) The evolutionary impact of invasive species. Proc Natl Acad Sci U S A, 98, 5446–5451.

Morrant, D.S., Turner, J.M., Jensen, M.A., Hansen, N.A., Bower, D.S., Körtner, G., Meek, P.D., Pestell, A.J., Rismiller, P.D. & Waudby, H.P. (2022) Wildlife tracking methods. Wildlife Research in Australia: Practical and Applied Methods, 180.

Morton, S.R. (1982) Dasyurid marsupials of the Australian arid zone: an ecological review. Carnivorous Marsupials (ed. M. Archer). Royal Zoological Society of New South Wales, Sydney, Australia.

Moseby, K.E., Hodgens, P., Bannister, H., Mooney, P., Brandle, R., Lynch, C., Young, C., Jansen, J. & Jensen, M. (2021a) The ecological costs and benefits of a feral cat poison-baiting programme for protection of reintroduced populations of the western quoll and brushtail possum. Austral Ecology, 46, 1366–1382.

Moseby, K.E., Hodgens, P., Peacock, D., Mooney, P., Brandle, R., Lynch, C., West, R., Young, C.M., Bannister, H., Copley, P. & Jensen, M.A. (2021b) Intensive monitoring, the key to identifying cat predation as a major threat to native carnivore (Dasyurus geoffroii) reintroduction. Biodiversity and Conservation, 30, 1547–1571.

Moseby, K.E., Jensen, M.A. & Tatler, J. (2022) Dietary flexibility and high predator efficacy facilitate coexistence in a novel predator interaction. Journal of Mammalogy, 103, 124–135.

Myers, K. (1970) The rabbit in Australia. Proceedings of the advanced study institute on dynamics of numbers in populations (eds P.J.d. Boer & G.R. Gradwell), pp. 478–506. Centre for Agricultural Publishing and Documentation,, Wageningen.

Palmas, P., Gouyet, R., Oedin, M., Millon, A., Cassan, J.J., Kowi, J., Bonnaud, E. & Vidal, E. (2020) Rapid recolonisation of feral cats following intensive culling in a semi-isolated context. NeoBiota, 63, 177–200.

Pavey, C.R., Addison, J., Brandle, R., Dickman, C.R., McDonald, P.J., Moseby, K.E. & Young, L.I. (2017) The role of refuges in the persistence of Australian dryland mammals. Biol Rev Camb Philos Soc, 92, 647–664.

Pearman, P.B., Guisan, A., Broennimann, O. & Randin, C.F. (2008) Niche dynamics in space and time. Trends Ecol Evol, 23, 149–158.

Pedler, R.D., Brandle, R., Read, J.L., Southgate, R., Bird, P. & Moseby, K.E. (2016) Rabbit biocontrol and landscape-scale recovery of threatened desert mammals. Conserv Biol, 30, 774–782.

Peers, M.J., Thornton, D.H. & Murray, D.L. (2013) Evidence for large-scale effects of competition: niche displacement in Canada lynx and bobcat. Proc Biol Sci, 280, 20132495.

Pringle, R.M., Kartzinel, T.R., Palmer, T.M., Thurman, T.J., Fox-Dobbs, K., Xu, C.C.Y., Hutchinson, M.C., Coverdale, T.C., Daskin, J.H., Evangelista, D.A., Gotanda, K.M., N, A.M.I.t.V., Wegener, J.E., Kolbe, J.J., Schoener, T.W., Spiller, D.A., Losos, J.B. & Barrett, R.D.H. (2019) Predator-induced collapse of niche structure and species coexistence. Nature, 570, 58–64.

QGIS Development Team (2020) QGIS Geographic Information System. Open Source Geospatial Foundation Project.

R Core Team (2020) R: A language and environment for statistical computing. R Foundation for Statistical Computing.

Richards, S.A. (2008) Dealing with overdispersed count data in applied ecology. Journal of Applied Ecology, 45, 218–227.

RStudio Team (2020) RStudio: Integrated Development Environment for R. RStudio, PBC.

Sale, P.F. (1974) Overlap in resource use, and interspecific competition. Oecologia, 17, 245–256.

SARIG (2021) South Australian Resources Information Gateway-100k Geology. Department for Energy and Mining, Government of South Australia.

Scheele, B.C., Foster, C.N., Banks, S.C. & Lindenmayer, D.B. (2017) Niche Contractions in Declining Species: Mechanisms and Consequences. Trends Ecol Evol, 32, 346–355.

Schoener, T.W. (1971) Theory of feeding strategies. Annual review of ecology and systematics, 2, 369–404.

Serena, M. & Soderquist, T.R. (1989) Spatial-Organization of a Riparian Population of the Carnivorous Marsupial Dasyurus-Geoffroii. Journal of Zoology, 219, 373–383.

Shelford, V.E. (1931) Some Concepts of Bioecology. Ecology, 12, 455–467.

Soderquist, T. & Serena, M. (1994) Dietary niche of the western quoll, Dasyurus geoffroii, in the jarrah forest of Western Australia. Australian Mammalogy, 17, 133–136.

Stead-Richardson, E., Bradshaw, S., Bradshaw, F. & Gaikhorst, G. (2001) Monitoring the oestrous cycle of the chuditch (Dasyurus geoffroii)(Marsupialia: Dasyuridae): non-invasive analysis of faecal oestradiol-17b. Australian Journal of Zoology, 49, 183–193.

Stevens, A.N.P. (2012) Dynamics of Predation. Papers in Natural Resources, 976.

Stobo-Wilson, A.M., Brandle, R., Johnson, C.N. & Jones, M.E. (2020) Management of invasive mesopredators in the Flinders Ranges, South Australia: effectiveness and implications. Wildlife Research, 47, 720–730.

U.S. Geological Survey (2022) Earth Explorer-Sentinel 2-USGS EROS Archive.

West, R.S., Tilley, L. & Moseby, K.E. (2020) A trial reintroduction of the western quoll to a fenced conservation reserve: implications of returning native predators. Australian Mammalogy, 42, 257–265.

Willoughby, N., Thompson, D., Royal, M. & Miles, M. (2018) South Australian land cover layers: an introduction and summary statistics. Department for Environment and Water, Adelaide.

Wilson, N., Tickle, P.K., Gallant, J., Dowling, T. & Read, A. (2011) 1 second SRTM Derived Hydrological Digital Elevation Model (DEM-H) version 1.0.

Woinarski, J.C., Burbidge, A.A. & Harrison, P.L. (2014) The action plan for Australian mammals 2012. CSIRO Publishing, Collingwood, Vic.

Woinarski, J.C., Burbidge, A.A. & Harrison, P.L. (2015) Ongoing unraveling of a continental fauna: decline and extinction of Australian mammals since European settlement. Proc Natl Acad Sci U S A, 112, 4531–4540.

