## Supplementary material for "Spatial niche separation of an invasive and a native mesopredator"

**S 1:** trap data of feral cats.

| ID | Sex | Weight in g | Collar % of Bodyweight | Days collared |
|---|---|---|---|---|
| C1120 | f | 3450 | 2.61 | 119 |
| C1720 | f | 2475 | 3.64 | 51 |
| C1820 | f | 1975 | 4.05 | 41 |
| C419 | f | 2170 | 3.69 | 36 |
| C519 | f | 2650 | 4.15 | 24 |
| C619 | f | 2450 | 3.27 | 50 |
| C719 | f | 2750 | 4 | 21 |
| C720 | f | 2025 | 3.95 | 10 |
| C820 | f | 2430 | 3.42 | 42 |
| C1019 | m | 2260 | 3.54 | 19 |
| C119 | m | 3950 | 2.03 | 48 |
| C120 | m | 2100 | 3.81 | 39 |
| C1220 | m | 4400 | 2.05 | 78 |
| C1319 | m | 2345 | 1.92 | 18 |
| C1420 | m | 2550 | 3.53 | 77 |
| C1520 | m | 2450 | 3.67 | 74 |
| C1620 | m | 2355 | 3.82 | 99 |
| C1920 | m | 4830 | 1.86 | 87 |
| C2020 | m | 2830 | 3.18 | 65 |
| C219 | m | 1600 | 5 | 41 |
| C220 | m | 2720 | 3.27 | 103 |
| C319 | m | 3400 | 2.35 | NA |
| C320 | m | 4300 | 2.09 | 11 |
| C420 | m | 2425 | 3.42 | 0 |
| C520 | m | 4500 | 1.98 | 23 |
| C620 | m | 4050 | 2.22 | 128 |
| C819 | m | 3750 | 2.93 | 20 |
| C919 | m | 3000 | 3.67 | 23 |
| C920 | m | 2080 | 4.28 | 19 |

**S 2:** trap data of western quolls

| ID | Sex | Weight in g | Collar % of Bodyweight | Days collared |
|---|---|---|---|---|
| Q1120 | f | 1080 | 2.78 | 19 |
| Q1520 | f | 1067.5 | 2.81 | 9 |
| Q1920 | f | 990 | 3.04 | 28 |
| Q2520 | f | 1000 | 3 | 24 |
| Q320 | f | 830 | 3.61 | 25 |
| Q619 | f | 985 | 2.84 | 55 |
| Q1019 | m | 1955 | 1.43 | 33 |
| Q1019_20 | m | 1725 | 1.74 | 24 |
| Q1220 | m | 1360 | 2.28 | 18 |
| Q1320 | m | 1475 | 2.03 | 28 |
| Q1620 | m | 1170 | 2.65 | 22 |
| Q1720 | m | 1360 | 2.35 | 18 |
| Q1820 | m | 1240 | 2.5 | 19 |
| Q2020 | m | 1480 | 2.16 | 21 |
| Q219 | m | 1210 | 2.31 | 56 |
| Q2220 | m | 1140 | 2.63 | 26 |
| Q2320 | m | 1405 | 2.14 | 27 |
| Q2420 | m | 930 | 3.23 | 16 |
| Q2620 | m | 1400 | 2.14 | 26 |
| Q2720 | m | 1680 | 1.79 | 24 |
| Q2820 | m | 1520 | 1.97 | 33 |
| Q2920 | m | 1535 | 1.95 | 32 |
| Q3020 | m | 990 | 3.03 | 46 |
| Q319 | m | 1960 | 1.43 | 65 |
| Q419 | m | 1670 | 1.68 | 74 |
| Q519 | m | 1300 | 2.15 | 55 |
| Q719 | m | 1590 | 1.76 | 71 |
| Q719_20 | m | 1675 | 1.79 | 49 |
| Q720 | m | 1110 | 2.7 | 32 |
| Q819 | m | 1020 | 2.75 | 47 |
| Q919 | m | 1310 | 2.14 | 22 |
| Q920 | m | 1250 | 2.4 | 29 |

**S 3:** movement and home range data of feral cats.

| ID | Nr of GPS fixes | mean distance in km/ day | sd distance | median distance in km/ day | mean speed in km/ h | sd speed | median speed in km/ h | MCP area in ha | Kref area in ha | BRB area in ha |
|---|---|---|---|---|---|---|---|---|---|---|
| C1120 | 5039 | 5.75 | 2.75 | 5.19 | 0.37 | 0.59 | 0.07 | 1,077 | 955 | 989 |
| C1720 | 3886 | 6.81 | 2.29 | 6.54 | 0.41 | 0.44 | 0.21 | 592 | 658 | 751 |
| C1820 | 2521 | 4.07 | 2.04 | 3.56 | 0.31 | 0.38 | 0.12 | 545 | 617 | 701 |
| C419 | 419 | 3.55 | 2.20 | 4.54 | 0.49 | 0.53 | 0.38 | 574 | 836 | 615 |
| C519 | 640 | 2.26 | 1.29 | 2.08 | 0.42 | 0.47 | 0.2 | 305 | 369 | 545 |
| C619 | 2930 | 4.41 | 2.89 | 3.53 | 0.31 | 0.68 | 0.07 | 1206 | 632 | 936 |
| C719 | 721 | 3.70 | 1.36 | 3.56 | 0.58 | 0.55 | 0.41 | 796 | 965 | 1,188 |
| C720 | 727 | 7.03 | 4.09 | 7.69 | 0.5 | 0.62 | 0.31 | 907 | 1244 | 923 |
| C820 | 2183 | 7.10 | 4.08 | 7.14 | 0.46 | 0.62 | 0.1 | 40,827 | 46,040 | 5,588 |
| C1019 | 1078 | 6.24 | 4.40 | 6.31 | 0.52 | 0.58 | 0.3 | 1,036 | 1,555 | 1,161 |
| C119 | 2401 | 6.30 | 2.28 | 6.80 | 0.54 | 0.52 | 0.42 | 671 | 753 | 1,027 |
| C120 | 1754 | 7.23 | 2.68 | 7.08 | 0.37 | 0.66 | 0.1 | 23,355 | 54,441 | 5,312 |
| C1220 | 6648 | 6.00 | 2.93 | 5.97 | 0.31 | 0.47 | 0.06 | 2,510 | 2,226 | 1,946 |
| C1319 | 518 | 3.15 | 1.67 | 2.75 | 0.56 | 0.54 | 0.32 | 1,885 | 4,763 | 1,130 |
| C1420 | 4553 | 4.01 | 2.36 | 3.69 | 0.28 | 0.38 | 0.09 | 793 | 555 | 866 |
| C1520 | 2233 | 5.05 | 3.20 | 4.50 | 0.32 | 0.72 | 0.1 | 893 | 757 | 800 |
| C1620 | 1638 | 2.75 | 1.98 | 2.55 | 0.24 | 0.38 | 0.04 | 729 | 769 | 823 |
| C1920 | 7600 | 13.38 | 4.09 | 14.40 | 0.71 | 0.91 | 0.17 | 5,143 | 3,347 | 3,387 |
| C2020 | 4869 | 5.02 | 2.66 | 4.36 | 0.3 | 0.46 | 0.05 | 966 | 1,514 | 1,338 |
| C219 | 1409 | 4.25 | 2.05 | 4.12 | 0.52 | 0.49 | 0.43 | 1,458 | 1,379 | 1,123 |
| C220 | 6390 | 6.07 | 3.07 | 6.01 | 0.38 | 0.56 | 0.11 | 56,804 | 47,856 | 5,840 |
| C319 | 1085 | 5.33 | 2.29 | 5.13 | 0.51 | 0.47 | 0.44 | 321 | 404 | 510 |
| C320 | 456 | 7.76 | 3.91 | 9.15 | 0.46 | 0.52 | 0.23 | 1,761 | 6,106 | 1113 |
| C420 | 727 | 7.03 | 4.09 | 7.69 | 0.5 | 0.62 | 0.31 | 907 | 1,244 | 923 |
| C520 | 1036 | 3.99 | 2.38 | 3.71 | 0.24 | 0.52 | 0.05 | 1,479 | 2,500 | 1,525 |
| C620 | 8056 | 8.88 | 4.30 | 8.27 | 0.47 | 0.66 | 0.08 | 157,834 | 213,051 | 12,533 |
| C819 | 864 | 3.14 | 1.84 | 2.85 | 0.34 | 0.39 | 0.17 | 1,184 | 1,451 | 1,130 |
| C919 | 848 | 3.60 | 1.74 | 3.95 | 0.39 | 0.39 | 0.21 | 881 | 1,127 | 815 |
| C920 | 104 | 5.87 | 4.12 | 4.22 | 0.77 | 0.63 | 0.75 | 1,013 | 2,438 | 731 |

**S 4:** movement data and home range sizes of western quolls.

| ID | Nr of GPS fixes | mean distance in km/ day | sd distance | median distance in km/ day | mean speed in km/ h | sd speed | median speed in km/ h | MCP area in ha | Kref area in ha | BRB area in ha |
|---|---|---|---|---|---|---|---|---|---|---|
| Q1120 | 570 | 2.41 | 1.80 | 1.81 | 0.33 | 0.52 | 0.07 | 335 | 384 | 478 |
| Q1520 | 558 | 1.59 | 0.97 | 1.44 | 0.25 | 0.41 | 0.06 | 95 | 111 | 392 |
| Q1920 | 437 | 1.50 | 1.17 | 1.19 | 0.38 | 0.54 | 0.12 | 250 | 389 | 637 |
| Q2520 | 580 | 4.34 | 2.83 | 4.39 | 0.71 | 0.75 | 0.47 | 421 | 531 | 655 |
| Q320 | 177 | 1.71 | 1.50 | 1.00 | 0.68 | 0.66 | 0.5 | 331 | 460 | 493 |
| Q619 | 729 | 2.58 | 1.62 | 2.40 | 0.44 | 0.52 | 0.23 | 531 | 665 | 705 |
| Q1019 | 891 | 5.15 | 3.91 | 4.42 | 0.86 | 1.02 | 0.37 | 6,318 | 10,278 | 4,602 |
| Q1019_20 | 519 | 8.52 | 4.72 | 7.23 | 1.25 | 1.5 | 1.04 | 6,965 | 10,206 | 5,694 |
| Q1220 | 217 | 4.65 | 3.89 | 3.58 | 1 | 0.89 | 0.94 | 2,113 | 3,278 | 1,752 |
| Q1320 | 283 | 3.49 | 2.22 | 3.27 | 0.63 | 0.73 | 0.24 | 1,832 | 3,695 | 1,644 |
| Q1620 | 54 | 3.33 | 4.25 | 1.84 | 1.57 | 1.05 | 1.73 | 2,504 | 5,384 | 1,986 |
| Q1720 | 331 | 4.67 | 2.87 | 4.59 | 0.77 | 1 | 0.21 | 1,555 | 2,372 | 1,679 |
| Q1820 | 159 | 4.29 | 1.86 | 4.08 | 0.95 | 0.87 | 0.85 | 2,840 | 7,833 | 2,640 |
| Q2020 | 53 | 5.42 | 4.12 | 5.42 | 1.08 | 1.26 | 0.29 | 1,854 | 6,276 | 900 |
| Q219 | 229 | 1.00 | 0.82 | 0.90 | 0.42 | 0.61 | 0.11 | 2,715 | 4,374 | 1,947 |
| Q2220 | 377 | 5.31 | 2.62 | 5.08 | 1.22 | 1.05 | 1.08 | 4,646 | 10,023 | 3,127 |
| Q2320 | 467 | 7.16 | 4.28 | 5.38 | 1.36 | 1.25 | 1.19 | 4,830 | 7,150 | 6,718 |
| Q2420 | 427 | 10.33 | 6.31 | 8.59 | 1.84 | 1.62 | 1.78 | 6,426 | 11,844 | 7,217 |
| Q2620 | 863 | 5.66 | 3.18 | 5.21 | 0.74 | 0.9 | 0.3 | 1,940 | 2,445 | 2,577 |
| Q2720 | 136 | 5.58 | 3.88 | 5.35 | 1.03 | 1.07 | 0.7 | 2,152 | 4,148 | 1,842 |
| Q2820 | 85 | 4.42 | 3.82 | 4.43 | 1.07 | 1.06 | 0.68 | 1,550 | 5,186 | 1,019 |
| Q2920 | 458 | 6.99 | 4.44 | 6.91 | 1.53 | 1.38 | 1.35 | 6,572 | 8,794 | 7,018 |
| Q3020 | 116 | 8.44 | 2.91 | 9.12 | 1.47 | 1.35 | 1.31 | 6,226 | 18,666 | 2,616 |
| Q319 | 786 | 5.67 | 2.63 | 5.37 | 0.86 | 0.84 | 0.57 | 6,101 | 8,043 | 4,228 |
| Q419 | 510 | 2.57 | 2.26 | 1.97 | 0.66 | 0.86 | 0.19 | 4,462 | 5,892 | 3,761 |
| Q519 | 579 | 1.34 | 1.31 | 0.93 | 0.34 | 0.66 | 0.06 | 587 | 1,063 | 1,247 |
| Q719 | 831 | 3.88 | 2.30 | 3.47 | 0.69 | 0.82 | 0.29 | 2,816 | 3,944 | 2,790 |
| Q719_20 | 510 | 5.63 | 3.91 | 4.69 | 0.99 | 1.04 | 0.61 | 6,530 | 7,752 | 4,380 |
| Q720 | 666 | 4.78 | 2.49 | 4.30 | 0.81 | 0.89 | 0.42 | 1,925 | 2,838 | 2,079 |
| Q819 | 83 | 2.76 | 2.30 | 2.31 | 0.93 | 0.79 | 0.79 | 911 | 2,757 | 744 |
| Q919 | 342 | 2.04 | 1.21 | 2.26 | 0.65 | 0.76 | 0.24 | 5,788 | 7,053 | 3,355 |
| Q920 | 493 | 6.80 | 4.58 | 5.78 | 1.12 | 0.97 | 1.03 | 1,401 | 1,715 | 2,060 |

**S 5:** measurements of precision of GPS/VHF collars of cats.

| ID | number of fixes | prec med | mean | sd | comment |
|---|---|---|---|---|---|
| C119 | 147 | 2.00 | 2.63 | 2.20 | |
| C120 | 548 | 8.97 | 14.77 | 17.92 | |
| C219 | 278 | 3.08 | 9.98 | 23.97 | |
| C320 | 176 | 9.06 | 14.51 | 17.23 | |
| C420 | 31 | 4.24 | 6.82 | 9.37 | |
| C520 | 637 | 5.83 | 13.44 | 40.05 | |
| C719 | 55 | 18.87 | 31.27 | 35.07 | |
| C720 | 31 | 4.24 | 6.82 | 9.37 | |
| C819 | 435 | 5.10 | 10.71 | 26.16 | |
| C820 | 683 | 8.49 | 13.01 | 15.31 | |
| C919 | 402 | 13.04 | 17.50 | 15.17 | |
| C920 | 1407 | 11.00 | 18.48 | 37.99 | |
| C1220 | 349 | 3.61 | 5.35 | 5.08 | |
| C1420 | 100 | 9.03 | 27.39 | 51.55 | |
| C1720 | 98 | 7.07 | 10.08 | 9.32 | |
| Cats median | 278.00 | 7.07 | 13.01 | 17.23 | |
| Cats mean | 358.47 | 7.57 | 13.52 | 21.05 | |
| Cats sd | 363.69 | 4.43 | 7.83 | 14.45 | |

**S 6:** measurements of precision of GPS/VHF collars of quolls.

| ID | number of fixes | prec med | mean | sd | comment |
|---|---|---|---|---|---|
| Q1120 | 162 | 8.54 | 18.55 | 39.69 | |
| Q1220 | 542 | 7.45 | 23.13 | 124.03 | |
| Q1320 | 793 | 8.06 | 25.19 | 145.20 | |
| Q1520 | 105 | 5.83 | 20.21 | 35.39 | |
| Q1620A | 633 | 8.49 | 22.46 | 194.69 | was moved |
| Q1620B | 398 | 10.31 | 21.41 | 34.44 | |
| Q1720 | 84 | 15.48 | 28.63 | 66.51 | |
| Q1820 | 617 | 6.00 | 15.35 | 46.53 | |
| Q2020 | 234 | 4.12 | 10.34 | 18.31 | |
| Q2420 | 109 | 11.40 | 19.07 | 21.13 | |
| Quolls median | 316.00 | 8.27 | 20.81 | 43.11 | |
| Quolls mean | 367.70 | 8.57 | 20.43 | 72.59 | |
| Quolls sd | 262.75 | 3.24 | 5.12 | 60.61 | |

**S7:** Summary of quoll distance models.

| model | df | AIC | delta_AIC | AIC_weight | Sexm | poly (day, 2)1 | poly (day, 2)2 | Sexm* poly (day, 2)1 | Sexm* poly (day, 2)2 | Weight |
|---|---|---|---|---|---|---|---|---|---|---|
| Sex * poly(day,2) + (1\|ID) | 8 | 6179.1 | 0 | 0.73 | 0.68 ± 0.14 | 3.40 ± 2.16 | 3.00 ± 1.44 | 3.97 ± 2.43 | -5.82 ± 1.63 | |
| Sex * poly(day,2) + Weight + (1\|ID) | 9 | 6181.1 | -1.98 | 0.27 | 0.66 ± 0.17 | 3.41 ± 2.16 | 3.01 ± 1.44 | 3.97 ± 2.42 | -5.82 ± 1.63 | 0.03 ± 0.22 |

**S8:** summary of the cat distance models.

| model | df | AIC | delta AIC | AIC weight | poly (day, 2)1 | poly (day, 2)2 | Weight | Sex |
|---|---|---|---|---|---|---|---|---|
| poly(day,2) + (1\|ID) | 5 | 17839.04 | 0 | 0.321076 | 1.41 ± 0.72 | 0.8 ± 0.67 | | |
| Weight + (1\|ID) | 4 | 17840.01 | -0.97447 | 0.197244 | | | 0.12 ± 0.07 | |
| Sex + (1\|ID) | 4 | 17842.83 | -3.79324 | 0.048185 | | | | 0.02 ± 0.13 |

**S9:** Home range analysis

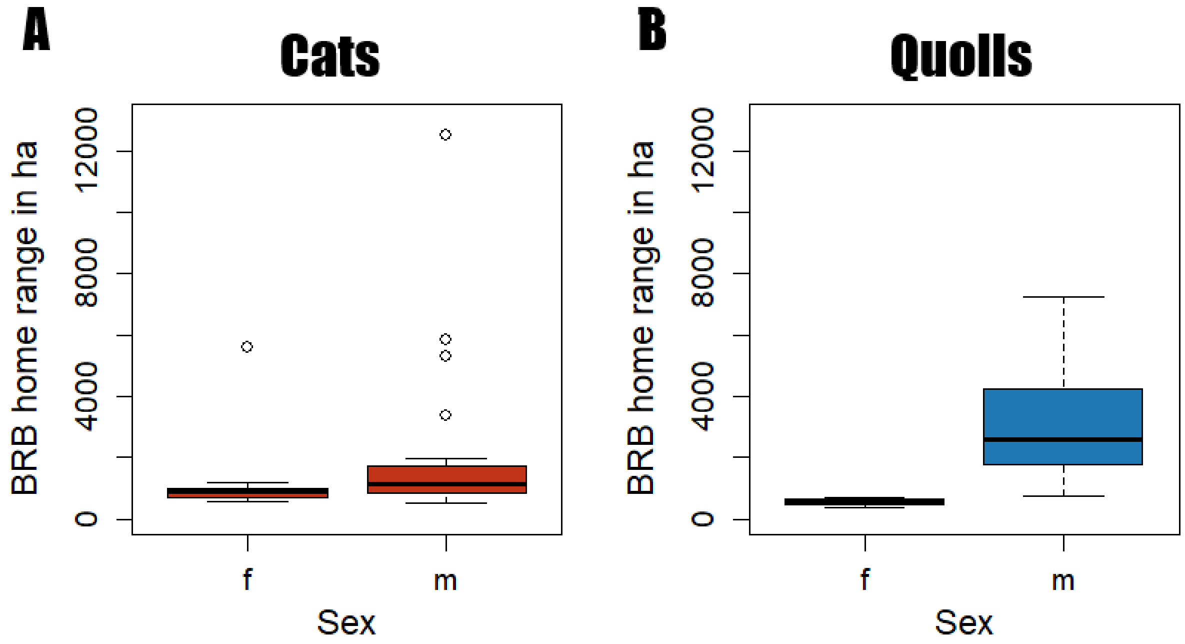

**S9.1:** 50% BRB home range sizes of A) female and male cats and B) female and male quolls.

**S9.2.: Welch Two Sample t-test**

**BRB home range size (**female and male quolls**)**

t = 6.70, df = 25.88, p-value = 4.21e-07

95% CI: 17.35; 32.70; mean of x: 30.62; mean of y: 5.60

**BRB home range size (**female and male cats**)**

t = 1.02, df = 25.36, p-value = 0.32

95% CI: -8.65; 25.49; mean of x: 22.02; mean of y: 13.60

**Daily distance (**female and male quolls**)**

t = 4.28, df = 16.57, p-value = 5.30e-04

95% CI: 1.34; 3.94; mean of x: 5.00; mean of y: 2.36

**Daily distance (**female and male cats**)**

t = 0.98, df = 21.15, p-value = 0.34

95% CI: -0.88; 2.46; mean of x: 5.76; mean of y: 4.96

**Speed t.test (**female and male quolls

t = 5.06, df = 14.82, p-value = 1.47 e-04

95% CI: 0.31, 0.75; mean of x: 0.99; mean of y: 0.47

**Speed t.test (**female and male cats**)**

t = 0.20, df = 24.02, p-value = 0.85

95% CI: -0.08; 0.10; mean of x: 0.44; mean of y: 0.43

**S 10: 50% BRB** home range overlap percentages of cats and quolls. Indicated are non-overlapping home ranges, and home ranges that overlap more than 5% of the area.

| Homerange of \ overlapped by | Cat | Quoll |
|---|---|---|
| **Cat** | No overlap: 92.4%<br>>5% overlap: 4.2% | No overlap: 95.6%<br>>5% overlap: 0.9% |
| **Quoll** | No overlap: 95.6%<br>>5% overlap: 0.5% | No overlap: 80.4%<br>>5% overlap: 8.5% |

**S 11: 99% BRB** home range overlap percentages of cats and quolls. Indicated are non-overlapping home ranges, and home ranges that overlap more than 5% of the area.

| Homerange of \ overlapped by | Cat | Quoll |
|---|---|---|
| **Cat** | No overlap: 74.9%<br>>5% overlap: 17.2% | No overlap: 74.1%<br>>5% overlap: 16.6% |
| **Quoll** | No overlap: 74.1%<br>>5% overlap: 12.3% | No overlap: 64.1%<br>>5% overlap: 30.4% |

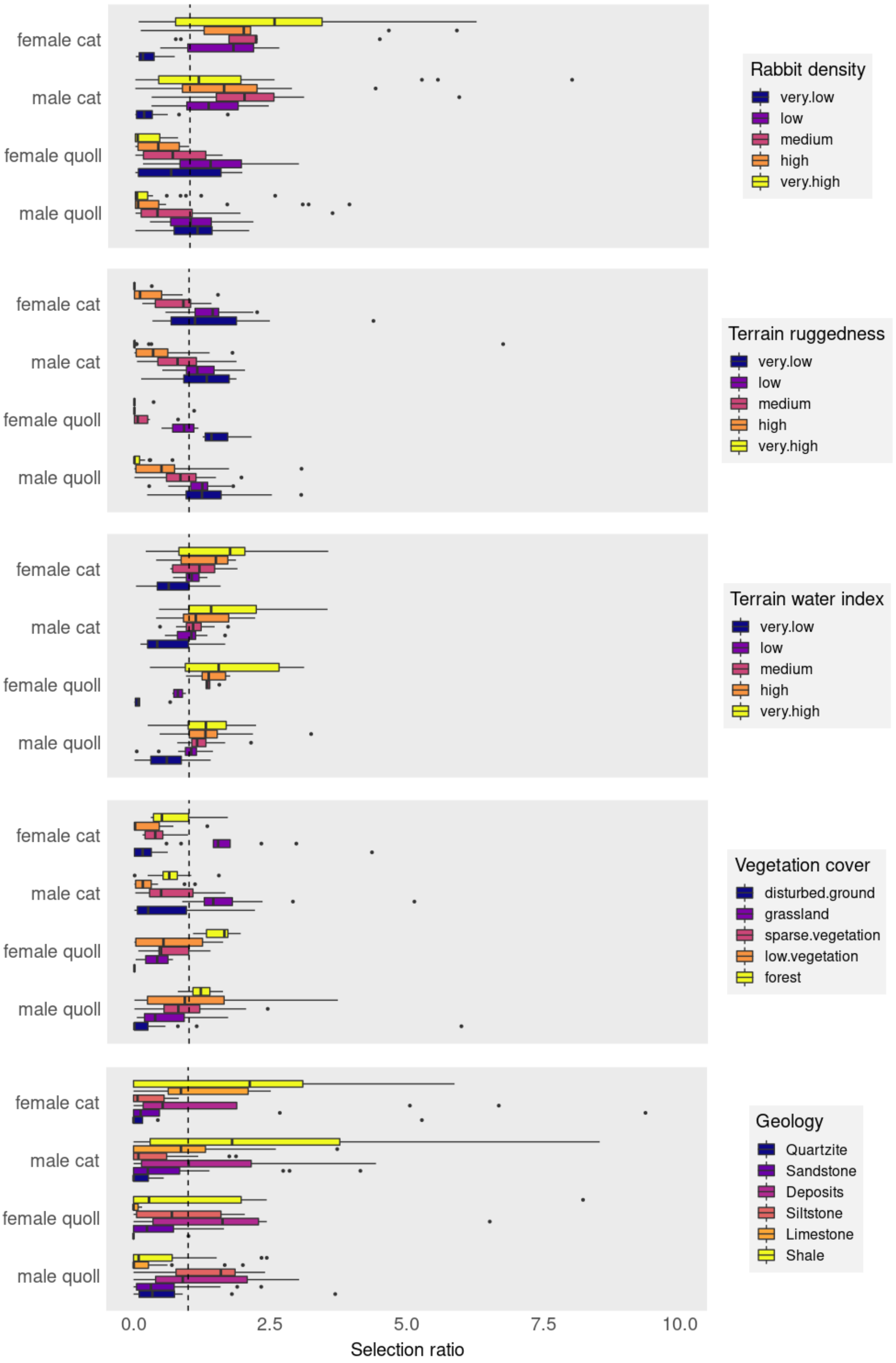

**S 12:** Habitat selection ratios of feral cats and western quolls within the Ikara-Flinders Ranges National Park and differences between different sexes for different environmental factors. Full plot including outliers. The dotted line represents no selection, values above are selected for, values below are selected against.

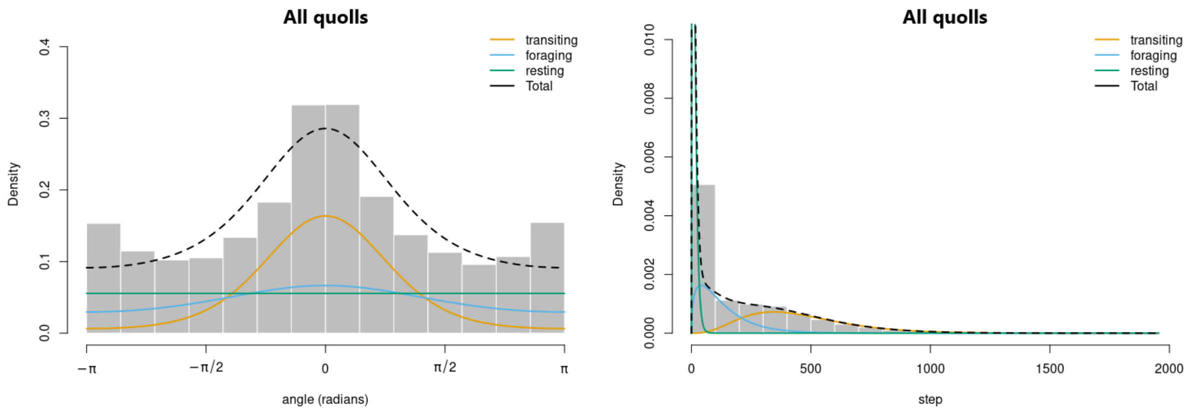

**S13:** HMM results of movement states of quolls.

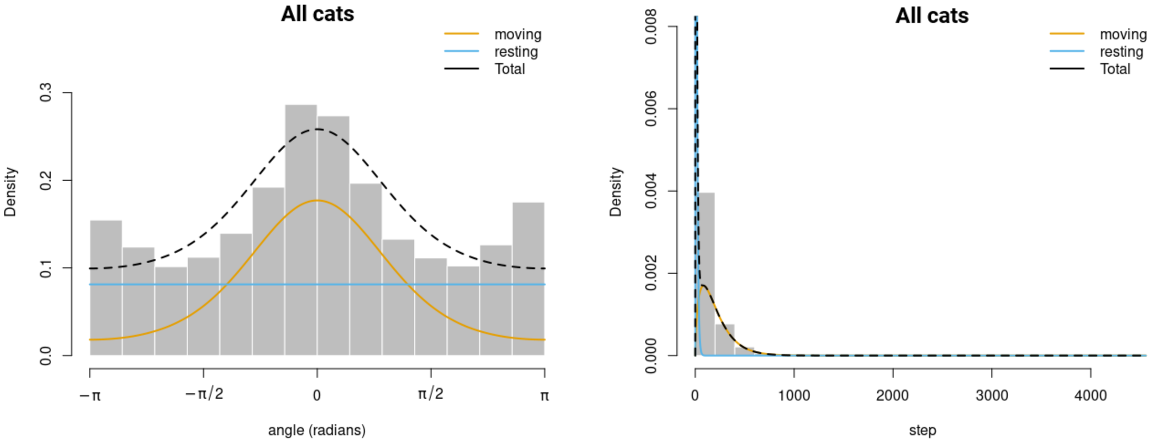

**S 14:** HMM results of movement states of cats.

**S 15:** results of the HMM models for movement states of female quolls.

| Model female quolls | AIC | delta_AIC | AIC_weights |
|---|---|---|---|
| PI+RABBITS | 37472.82 | 0 | 0.51 |
| PI | 37474 | -1.17 | 0.28 |
| PI+TRI | 37474.8 | -1.97 | 0.19 |
| PI+RABBITS+TRI | 37478.78 | -5.96 | 0.03 |

**S 16:** results of the HMM models for movement states of male quolls.

| Model male quolls | AIC | delta_AIC | AIC_weights |
|---|---|---|---|
| TRI | 137598.1 | 0 | 0.31 |
| TRI+PI | 137598.8 | -0.74 | 0.22 |
| TRI+NDVI | 137599.7 | -1.62 | 0.14 |
| 3 state (null) | 137600.3 | -2.21 | 0.10 |
| RABBITS | 137601.1 | -3.03 | 0.07 |
| PI | 137601.7 | -3.62 | 0.05 |
| NDVI+RABBITS | 137602.3 | -4.19 | 0.04 |
| NDVI | 137603.1 | -4.99 | 0.03 |
| TRI+RABBITS | 137603.3 | -5.22 | 0.02 |
| PI+RABBITS | 137604 | -5.89 | 0.02 |

**S 17:** results of the HMM models for movement states of female cats.

| Model female cat | AIC | delta_AIC | AIC_weights |
|---|---|---|---|
| GEO+ TRI | 239192.3 | 0 | 0.66 |
| GEO | 239194.7 | -2.38 | 0.20 |
| GEO+ TWI | 239197.4 | -5.13 | 0.05 |
| GEO+ PI | 239197.5 | -5.14 | 0.05 |
| GEO+ RABBITS | 239198.1 | -5.79 | 0.037 |

**S 18:** results of the HMM models for movement states of male cats.

| Model male cat | AIC | delta_AIC | AIC_weights |
|---|---|---|---|
| PI+ GEO | 701160.2 | 0 | 0.52 |
| PI+ GEO+ TRI | 701161.7 | -1.45 | 0.25 |
| PI+ TRI | 701162.2 | -1.94 | 0.20 |
| PI | 701165.5 | -5.31 | 0.04 |

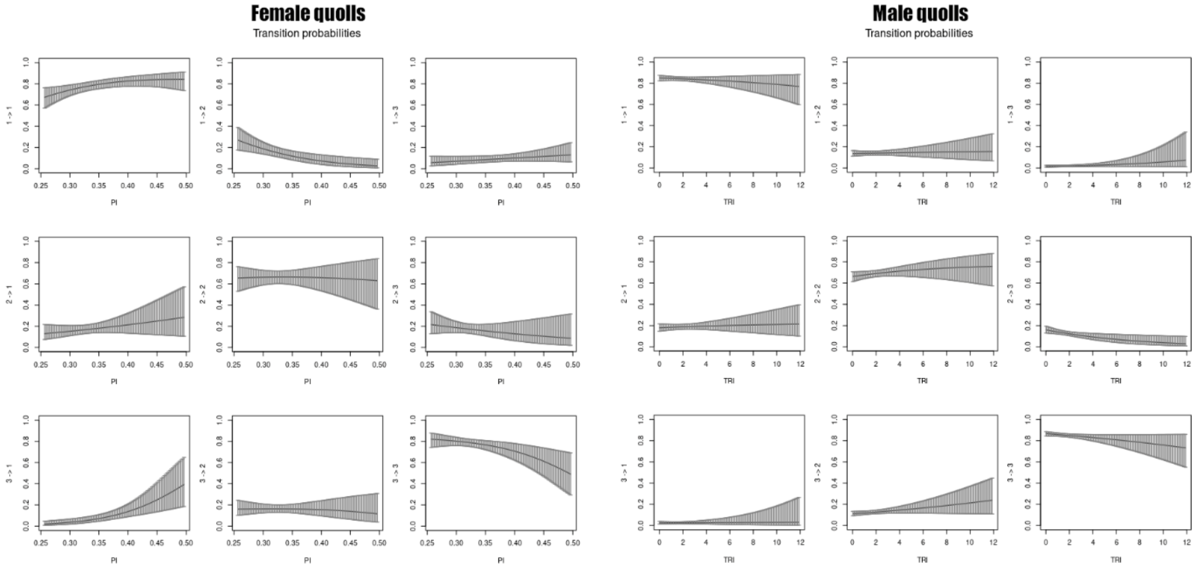

**S 19:** transition probabilities between 3 movement states calculated from the best model for female (left) and male quolls (right). Movement states are: 1= transiting; 2= foraging; 3= stationary.

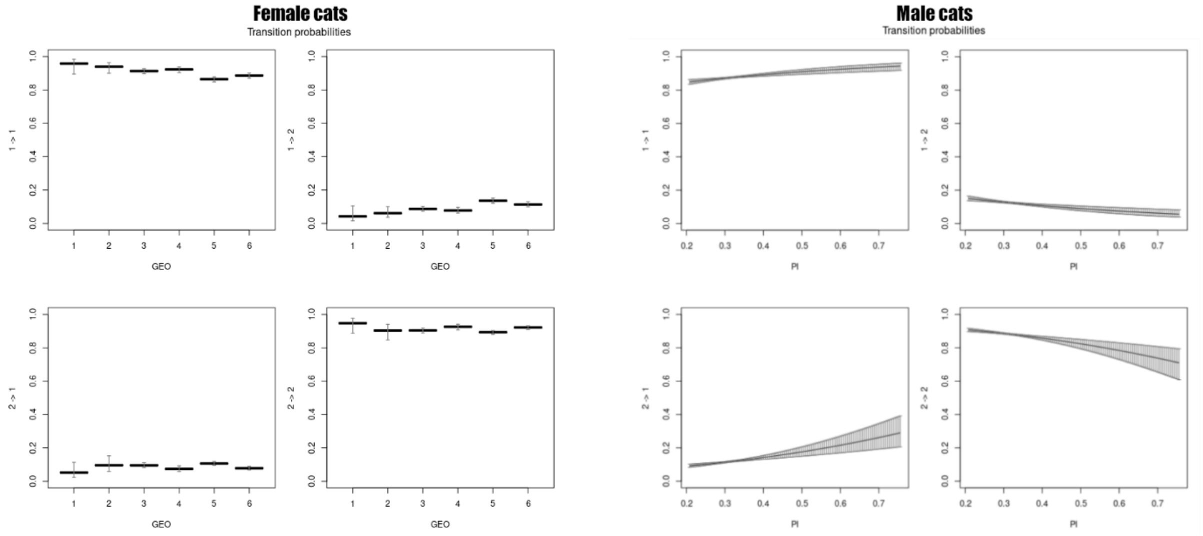

**S 20:** transition probabilities between 3 movement states calculated from the best model for female (left) and male cats (right). Movement states are: 1= transiting; 2= stationary.
